# Museum phylogenomics of extinct *Oryctes* beetles from the Mascarene Islands

**DOI:** 10.1101/2020.02.19.954339

**Authors:** Sergio M. Latorre, Matthias Herrmann, M.J. Paulsen, Christian Rödelsperger, Andreea Dréau, Waltraud Röseler, Ralf J. Sommer, Hernán A. Burbano

## Abstract

The evolution of island systems is characterized by processes that result in extreme morphological diversity, high endemism and high extinction rates. These dynamics can make phylogenetic reconstruction difficult, i.e. the extinct flightless Dodo from Mauritius was assigned to the family of doves only through DNA analysis of subfossils. Many insect species on islands have gone extinct through habitat loss, and face similar challenges to decipher their evolutionary history, however historical specimens have not yet been harnessed for phylogenomic reconstructions. Here, we employed historical museum specimens from the Mascarene Islands to generate the first whole-genome based phylogeny of three presumably extinct species of the rhinoceros beetle genus *Oryctes*. We compared their genomes with those of an extant *Oryctes* species from the island of Réunion, as well as a flightless Réunion-based species previously placed into the supposedly unrelated genus *Marronus*. We found that *Marronus borbonicus* belongs instead to the genus *Oryctes* and that the two Réunion-based species (*O. borbonicus* and *M. borbonicus*) are not sister taxa, suggesting two independent colonizations. The divergence time between them (<3Myr) overlaps with the volcanic formation of Réunion, likely indicating that *M. borbonicus* became flightless *in situ*. Our study showcases the power of genomes from insect museum specimens to address evolutionary questions in light of increasing extinction rates.

## Main

Coleoptera (beetles) are the most diverse order in metazoans with almost 400,000 described species^1^. Some lineages have given rise to spectacular forms that have fascinated humans for millennia. For example, illustrations featuring rhinoceros beetles have been found in Crete from the Minoan period (2000-1600 BC)^2^. Large and comprehensive museum collections of insect specimens exist throughout the world and the overall beetle phylogeny has been studied in great detail using both morphological and molecular tools based on a handful of loci^3,4^. The rhinoceros beetle genus *Oryctes* includes some of the largest beetles, such as *O. gigas*^*5*^, a species well known for its impressive horns^6^. In total, *Oryctes* contains 42 valid species distributed in Africa, Southeast Asia and the Indian Ocean^5^. The genus displays extremely high levels of endemism on islands in the Indian Ocean, which are considered major terrestrial biodiversity hotspots^7^. For instance, Madagascar and the Mascarene Islands (Réunion, Mauritius and Rodrigues) alone harbor 16 species, some of which have gone extinct due to extreme habitat loss, i.e. *O. tarandus* and *O. chevrolatii* on Mauritius and *O. minor* on Rodrigues (Figure 1A). While several systematic classifications have been suggested based on morphological characters, no molecular phylogeny is available because traditionally used PCR-based methods are not suitable for highly-degraded DNA present in historical museum specimens, and state-of-the-art library-based methods have not been employed for phylogenomic studies in extinct insect species. Given the distribution and diversity of rhinoceros beetles across the Mascarene Islands, establishing their evolutionary relationships is fundamental to understanding how geological processes, such as landmass emergence and submergence, have shaped the Mascarene’s endemic biodiversity.

**Figure 1.**
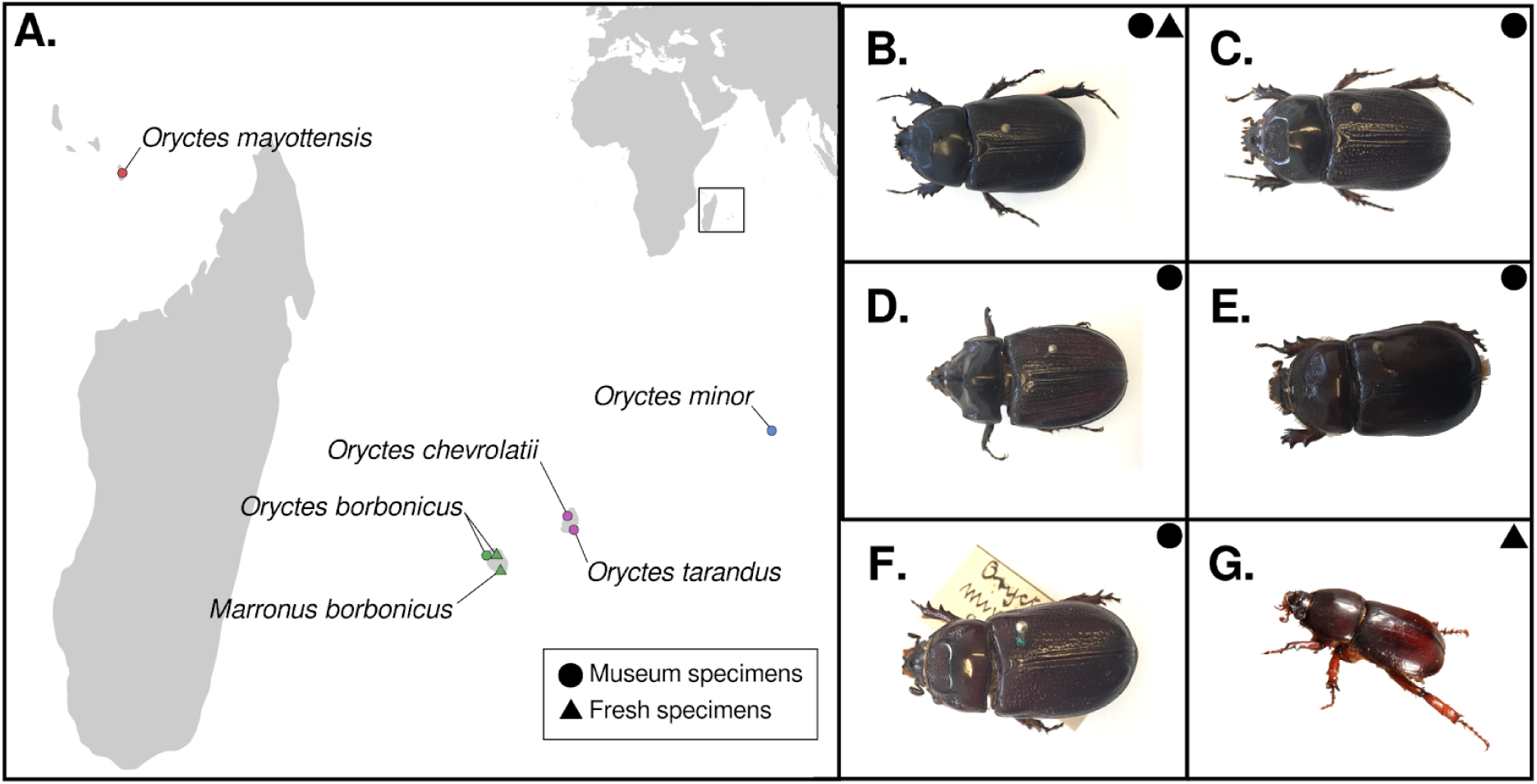
Provenance of fresh and museum rhinoceros beetle specimens. **(A)** The map displays beetle species endemic to different islands in the Indian Ocean on the coast of East Africa. The largest island is Madagascar. From west to east the sampled species included *Oryctes mayottensis* in Mayotte, *Oryctes borbonicus* and *Marronus borbonicus* in Réunion, *Oryctes chevrolatii* and *Oryctes tarandus* in Mauritius, and *Oryctes minor* in Rodrigues. Réunion, Mauritius and Rodrigues comprise the Mascarene Archipelago. *O. chevrolatii, O. tarandus* and *O. minor* have not been observed in decades and are presumably extinct. **(B-F)** Beetle museum specimens used in this study. **(B)** *O. borbonicus* **(C)** *O. tarandus* **(D)** *O. chevrolatii* **(E)** *O. minor* **(F)** *O. mayottensis* **(G)** Fresh specimen of *M. borbonicus*.

Réunion, the youngest of the Mascarene islands, harbors *O. borbonicus*, and an additional rhinoceros beetle that was placed in the monotypic genus *Marronus* based on morphological analyses^8^. This genus has been included in the tribe Pentodontini, and presumed to be distantly related to *Oryctes* in the tribe Oryctini. Like many island beetles, *M. borbonicus* (hereafter referred to as *Marronus* to avoid confusion with *Oryctes borbonicus*) is a flightless species that has undergone dwarfism. The creation of the monotypic genus *Marronus* based only on morphology is problematic since morphological features are often lost, and frequently convergent, in dwarf species. Moreover, the high prevalence of character displacement in islands can obscure morphological synapomorphies. Both *O. borbonicus* and *Marronus* are hosts of the nematode *Pristioncus pacificus*, a well-studied model organism for integrative evolutionary biology^9^. Thus, understanding the phylogenetic relationship between these two sympatric species might also shed light on the evolutionary history of the association between Réunion-based beetles and *P. pacificus*.

To investigate the phylogenetic relationships among *Oryctes* species from the Mascarene Islands and between them and *Marronus*, we used Illumina sequencing and 10X Genomics to refine the draft genome of *O. borbonicus*, and generated a new draft genome of *Marronus*. Furthermore, we used minute amounts of tissue from pinned beetle museum specimens to generate for the first time genome-wide data from extinct insect species from Mauritius and Rodrigues. The combined analysis of extant and extinct genomes permitted us to infer phylogenetic relationships, divergence times, and the colonization history of *Oryctes* beetles of the Mascarene Islands.

### *De novo* assembly of Reunion’s *Oryctes borbonicus* and *Marronus borbonicus*

Despite the large number of beetle species (∼400,000), genome sequences of only 20 species have been reported^10,11^. Available genomic data includes a draft assembly of *O. borbonicus*^12^. *Here, we sequenced DNA from two specimens of O. borbonicus* and *Marronus*, the two extant endemic beetles from Réunion, on the 10X Genomics platform to improve the draft genome of *O. borbonicus*, and to generate a draft genome of *Marronus*. Both libraries were individually sequenced on the Illumina HiSeq 3000 platform yielding 330 million paired end reads (2×150bp), which translates into roughly 120X coverage per genome. These data were assembled into 411 Mb for *O. borbonicus* and 413 Mb for *Marronus* (Supplementary Table 1). In comparison to the previously published draft genome of *O. borbonicus*^12^, this led to a huge reduction in contig number from over 150,000 to 9,526 and an 80-fold increased contiguity, i.e. N50 was raised from 105kb to 8.4Mb (Supplementary Table 1). Coverage analysis of the previous *O. borbonicus* assembly, revealed a high abundance of genomic regions with half the expected coverage possibly pointing towards a problem of allelism, i.e. two divergent haplotypes were assembled separately^12,13^. Indeed, sequencing of linked reads allowed the new *O. borbonicus* assembly to be resolved into pseudohaplotypes^14^. Consequently, the coverage profile of the new assembly is substantially shifted towards higher coverage, most likely autosomal regions (Supplementary Figure 1) indicating that the increased size of the previous assembly was largely due to separate assembly of divergent haplotypes. In total, 17,736 and 14,738 protein coding genes were annotated in the assemblies of *O. borbonicus* and *Marronus*, respectively. Comparison of protein sequence identity between 9,656 orthologous pairs revealed a median percentage identity of 98%, indicating that *O. borbonicus* and *Marronus* are closely related. Indeed, further phylogenomic analysis in the context of 13 phylogenetically broad Coleopteran genomes indicated that *O. borbonicus* and *Marronus* are closely related, similar in phylogenetic distance to that between *Hycleus phaleratus* and *Hycleus cichorii*, two members of the same genus (Supplementary Figure 2).

### Sampling and sequencing of historical beetle genomes

As three other endemic *Oryctes* species from both Mauritius (*O. tarandus* and *O. chevrolatii*) and Rodrigues (*O. minor*) are extinct, the only way to robustly reconstruct their evolutionary relationships is by retrieving genomes from historical museum specimens. To this purpose we obtained pinned museum specimens from these three extinct species, and also from *O. mayottensis* from Mayotte Island (as an outgroup for the Mascarene species), and an *O. borbonicus* museum specimen for comparison (Figure 1 and Supplementary Table 2). The age range of the museum specimens was between 53 and 99 years old. Although museum specimens from insects have been used before for phylogenetic analyses, this was mainly by PCR-based methods that aimed at amplifying one or a handful of loci^15,16^. In general, PCR amplifications have failed when using samples older than 50 years^17^. In contrast, whole-genome data derived from beetle museum specimens have not been used for phylogenetic analyses. To minimize the degree of sample destruction, we extracted DNA from one leg from each museum specimen amounting to ∼7-23 mg of tissue (Supplementary Figure 3A-E). The DNA extractions^18^ and library preparations were carried out in a clean room facility to avoid contamination from exogenous DNA. Shallow sequencing of the libraries showed damage patterns and length distributions typical of ancient DNA (aDNA) (Supplementary Figure 3F-G)^19^, and endogenous beetle DNA percentage varied from 5 to 85% (Supplementary Figure 3H). The variation in the percentage of endogenous DNA did not correlate with the distance of each species to the draft genomes of either *O. borbonicus* or *Marronus* (Supplementary Figure 3I). The same DNA extracts were used to generate chemically repaired libraries with significantly reduced ancient DNA-associated damage^20^ (Supplementary Figure 4), which were sequenced using the Illumina platform achieving on average 1X coverage (Supplementary Table 3). Only sequencing data derived from these chemical repaired libraries were used for subsequent analysis.

### Evolutionary relationships and divergence times of *Oryctes* spp. and *Marronus borbonicus*

To establish the evolutionary relationship among all beetle species, we first mapped the sequenced reads to the updated *O. borbonicus* draft genome, which provides a common coordinate system for all beetle species. To ascertain single nucleotide polymorphisms (SNPs) for each sample, we randomly selected one base at each segregating site. This method, also known as “pseudo-haploidization”, is commonly applied to low-coverage aDNA datasets, allowing the estimation of genetic relatedness from low-coverage data^21^ (See Materials and Methods). Implementing this methodology, we identified a total of 1,541,675 SNPs.

Initially, we summarized the genetic variation among beetle species using principal components analysis (PCA), which showed that PC1 does not separate species by genera but, instead, clusters together *Oryctes* and *Marronus* beetles from Réunion and Mauritius, and separates the species from Mayotte and Rodrigues (*O. mayottensis* and *O. minor*) from each other, and from the species from Réunion and Mauritius (Supplementary Figure 5A). This separation suggests that the vast majority of SNPs might have occurred in the lineages leading to *O. mayottensis* and *O. minor*. To test this hypothesis, we implemented a Minor Allele Frequency (MAF) filter that requires the SNPs to be segregating in at least two out of the seven samples (2/7). This filtering step reduced both the number of SNPs to 304,417, and the percentage of variance explained by PC2 in 30%. Indirectly, this reduction along the PC2 axis increased the separation among the beetles from Réunion and Mauritius (Figure 2A and Supplementary Figure 5). These observations support the hypothesis that the variance explained by PC2 was driven by variation that is private to either *O. mayottensis* or *O. minor* (Supplementary Figure 5A). Thus, after implementing the MAF filtering, the PC2 further separated beetle species. Importantly, PC2 also grouped together *O. borbonicus* from fresh and museum samples, which demonstrates that the use of chemically-repaired libraries and the appropriate identification of SNPs permit the accurate clustering of species, independent of their present-day or historical origin (Figure 2A and Supplementary Figure 6).

**Figure 2.**
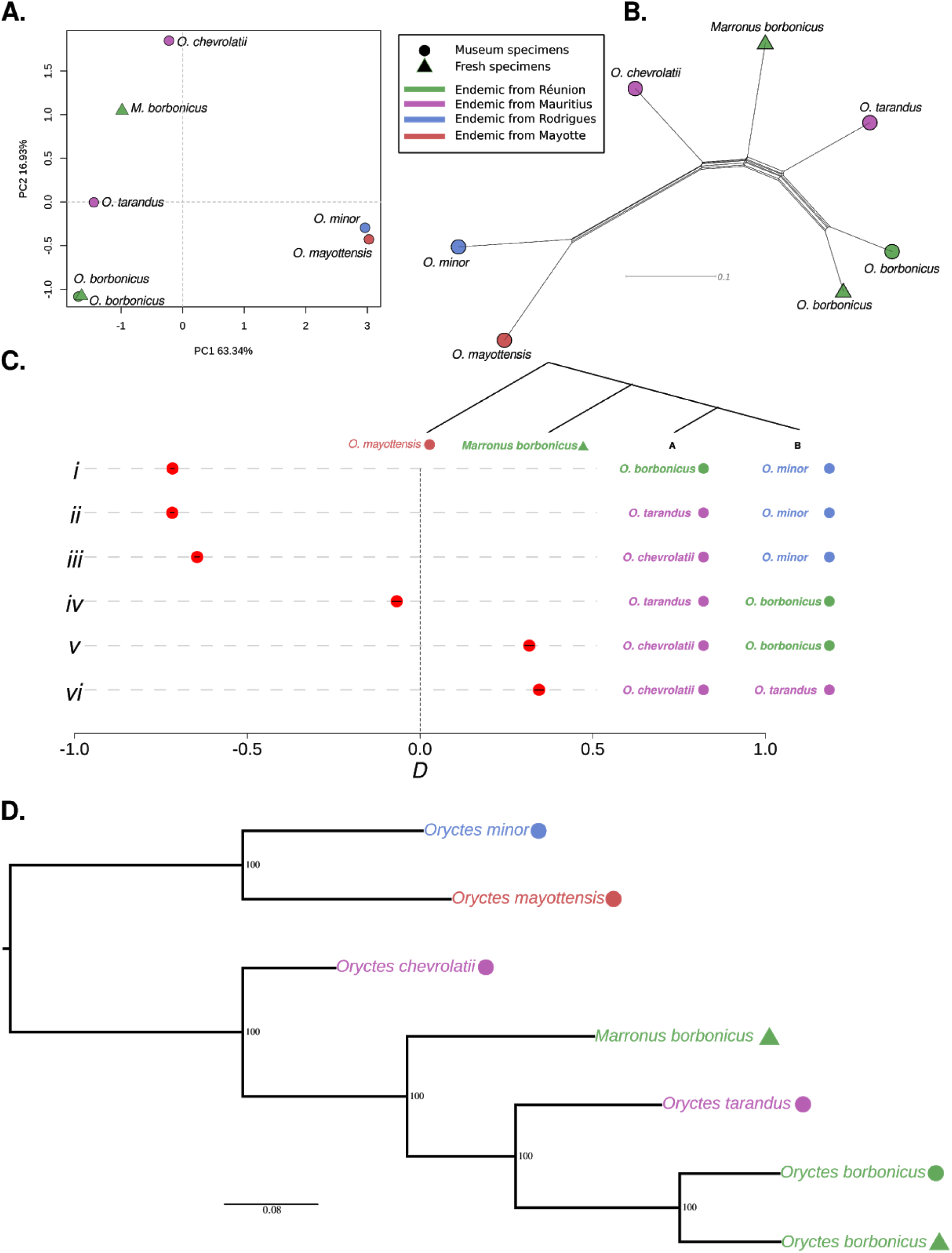
Evolutionary relations among rhinoceros beetles. **(A)** Principal component analysis plot based on 304,417 SNPs. Genetic distances between beetle samples are projected onto the first two PCs. Axis labels indicate the fraction of total variation explained by each PC. **(B)** Phylogenetic network based on 304,417 SNPs using the neighbor-net method. **(C)** Testing the robustness of phylogenetic relations among scarab beetle species using *D*-statistics of the type *D*(B,A; *Marronus borbonicus*, outgroup), as depicted in the phylogenetic tree. *O. mayottensis* was used as an outgroup. Each row (*i-vi*) shows a different *D*-statistic configuration. A negative *D*-statistic indicates that *M. borbonicus* is closer to species A, whereas a positive *D*-statistic indicates that *M. borbonicus* is closer to species B. The points depict the result of each *D*-statistic test and the lines their respective 95% confidence intervals. Rows *i-iii* show that *M. borbonicus* is closer to the *Oryctes* spp. from Réunion and Mauritius. Rows *v-vi* show that *M. borbonicus* is closer to both *O. borbonicus* and *O. tarandus* than to *O. chevrolatii*. Finally, row *iv* shows the closest *D*-statistic to zero, which indicates that *M. borbonicus* is slightly closer to *O. tarandus* than to *O. borbonicus* and **(D)** Maximum-Likelihood phylogenetic tree. Numbers at nodes indicate bootstrap support (200 replicates).

To further refine the evolutionary relationships among these Mascarene beetles, we built phylogenetic networks using either of the two SNP sets, i.e. with and without MAF filtering. As expected from the PCA analysis, the use of the MAF filtering reduced the branch lengths of *O. mayottensis* and *O. minor*, while preserving the network topology (Figure 2B and Supplementary Figure 6A-B). To focus on the evolutionary relationships among Réunion and Mauritius beetles, from here on, we carried out all analyses implementing the MAF 2/7 filtering. The phylogenetic networks revealed that (i) sympatric species pairs from Réunion and Mauritius do not cluster together but, instead, *O. borbonicus* and *O. tarandus* appear as sister groups, (ii) *Marronus* falls within the *Oryctes* genus, and (iii) *O. mayottensis* and *O. minor*, as suggested by the PCA analysis, are outgroups for beetles from Réunion and Mauritius. The absence of pervasive reticulation in the phylogenetic network might suggest that introgression is not substantial between Mascarene beetles.

In order to test the phylogenetic relationships (the “treeness”) suggested by the phylogenetic network, we used *D*-statistics^21,22^. We employed *D*-statistics of the following form: *D*(Outgroup, *Marronus*; species A, species B), using *O. mayottensis* as an outgroup and different configurations of the four *Oryctes* species from Réunion and Mauritius as species A and B (Figure 2C). The first three rows (*i-iii*) of Figure 2C indicated that *Marronus* is closer to *Oryctes* species from Reunión and Mauritius than to *O. minor* from Rodrigues, whereas the last two rows (*v-vi*) showed that *Marronus* is closer to both *O. borbonicus* and *O. tarandus* than to *O. chevrolatii*. The extreme negative and positive *D*-statistics of rows *i-ii* and *v-vi* indicated that the tested phylogenetic hypotheses are likely incorrect. Finally, row *iv*, where the *D*-statistic is the closest to zero, showed that *Marronus* is slightly closer to *O. tarandus* than to *O. borbonicus*, which could be explained by post-speciation introgression between *Marronus* and *O. tarandus*. To evaluate the influence of genome reference bias in our phylogenomic inferences, we repeated the *D*-statistic analyses but instead of mapping the reads to the *O. borbonicus* draft genome, we mapped the reads to the *Marronus* genome. *D*-statistics values for configurations presented in all rows but row *iv* showed qualitatively similar results, very negative for rows *i-iii* and very positive for rows *v-vi* (Figure 2C and Supplementary Figure 8C). Also consistent with previous analysis, row *iv* had the *D*-statistics closest to zero but this time with a positive value, which indicates a closer relationship between *Marronus* and *O. borbonicus* that again could be caused by post-speciation introgression, this time between *Marronus* and *O. borbonicus*. The fact that the sign of the *D*-statistic in row *iv* switched between negative and positive depending on the reference genome used very likely indicates that *Marronus* is equally distant to both *O. borbonicus* and *O. tarandus* and that the true value would overlap zero. A *D*-statistic not different from zero likely suggests that negligible or no post-speciation introgression took place between *Marronus* and either *O. borbonicus* and *O. tarandus*, thus any segment of the genome showing a discordant phylogeny is most likely the result of incomplete lineage sorting^21^. The lack of pervasive introgression suggested by both the lack of reticulations in the phylogenetic network and the *D*-statistics prompted us to carry out a phylogenetic analysis using genome-wide SNPs. Both Maximum Likelihood and Bayesian trees (Figure 2D and Supplementary Figure 7) confirmed that *Marronus* indeed belongs to the genus *Oryctes* and, thus, do not warrant a monotypic genus. Additionally, the fact that *O. borbonicus* and *O. tarandus*, although endemic to different islands, are sister groups, and that *Marronus* is an outgroup to both of them, likely indicates that Réunion was colonized independently by *O. borbonicus* and *Marronus*. All our results (Figure 2) proved to be robust to the choice of reference genome (Supplementary Figure 8).

To relate the speciation events with the geological processes that gave rise to the Mascarene Islands, we set out to calculate the divergence times between Mascarene’s beetles. Initially, to investigate the variation of DNA sequence divergence along the genome, we analyzed the nucleotide divergence of all species relative to the *O. borbonicus* draft genome using 100 kb windows (Fig 3A). As suggested by their phylogenetic relationship as outgroups for Réunion and Mauritius beetles, *O. minor* and *O. mayottensis* displayed the highest nucleotide divergence relative *O. borbonicus*, with medians of 3 and 3.8%, respectively. The use of pairwise genetic differences reflected the true genetic distance between *O. borbonicus* and both *O. mayottensis* and *O. minor* as shown in the non-MAF filtered genetic network (Supplementary Figure 6A). The distributions of nucleotide divergences between *O. borbonicus* and Réunion (*Marronus*) and Mauritius (*O. tarandus* and *O. chevrolatii*) beetles showed a high degree of overlap (Figure 3A), reflecting very close divergence times, as expected given their close phylogenetic relationships. Although, we found that mean pairwise nucleotide divergence does not perfectly correlate with the inferred phylogenetic relationships between beetle species, we expect that mean sequence divergence will provide a rough estimate of sequence divergence times. To translate mean sequence divergence between all sequenced beetle species and *O. borbonicus*, we used the most commonly used insect mutation rate: 1.15% (1.1%-1.2%) per million years^23^. Although this rate was calculated using mitochondrial DNA (mtDNA) from a different insect order (Lepidoptera)^23^, a recent attempt to calibrate the insect molecular clock using beetle mtDNA estimated a rate very similar to the Lepidoptera-based one^24^. Moreover, it has been suggested that mtDNA and nuclear mutation rates are very similar in insects^24,25^.

**Figure 3.**
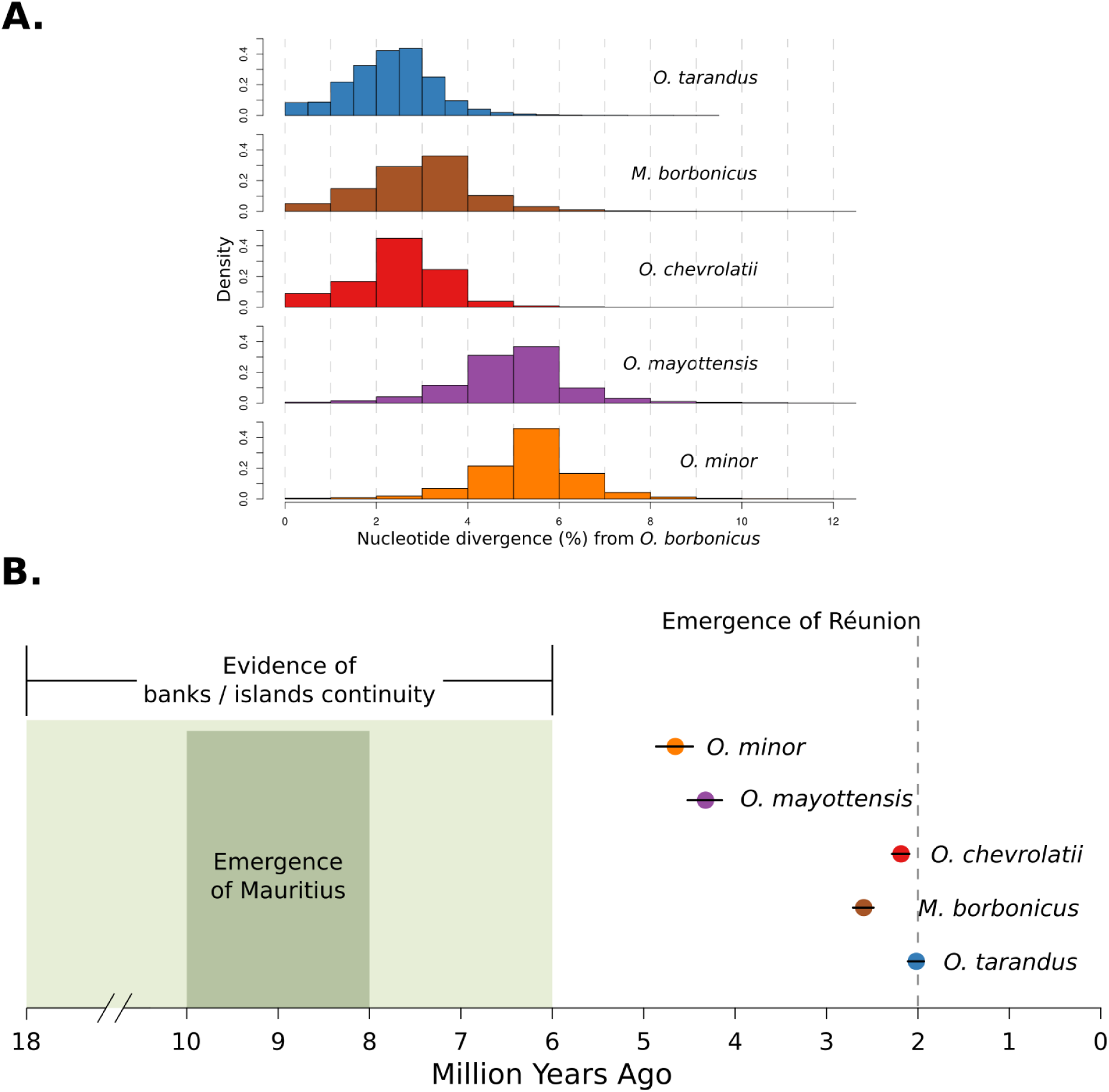
Nucleotide divergence of rhinoceros beetles. **(A)** Distribution of nucleotide divergence from the *Oryctes borbonicus* genome among genomic segments of 100 kb (N=1,553). **(B)** Divergence times of rhinoceros beetles relative to the *O. borbonicus* genome. The distributions of nucleotide divergences from (A) were converted into divergence times in million years using a constant substitution rate of 1.15 (1.1 - 1.2) % pairwise sequence divergence per million years per lineage^23^. The horizontal bars indicate upper and lower bounds for the divergence times based on the confidence intervals of the substitution rate. The x-axis shows the timing of major geological events in the Mascarene plateau, including the emergence of present-day Mascarene Islands.

The translation of mean sequence divergence resulted in sequence divergence times (in million years) of 1.70 (1.63-1.68) for *O. chevrolatii*, 1.90 (1.82-1.99) for *O. tarandus* and 1.79 (1.77-1.87) for *Marronus* (Figure 3B). Thus, all sequence divergence times between Réunion and Mauritius beetles relative to *O. borbonicus* are younger than 2 million years, and close in time to each other, showing overlapping confidence intervals in the case of *Marronus* and *O. tarandus*. Note that our approach to calculate sequence divergence times overestimates population divergence times, since sequence divergence times include the coalescence within the common ancestor of two species, while the population divergence - the point at which species stop exchanging genes - takes place much later in time. However, our estimated divergence times provide approximate estimates that can be overlaid with the geology of the Mascarene Islands. This overlay suggests that *Marronus* diverged from the common ancestor of *O. borbonicus* and *O. tarandus* at a time posterior to the emergence of Réunion (Figure 3B). These results imply that *Marronus* became flightless *in situ*, since it is less likely for a flightless species to be able to colonize an island, as has been proposed for the flightless Rodrigues solitaire^26^.

### Morphological analysis of *Marronus borbonicus* and its relation to *Oryctes* spp

In light of the phylogenomic analyses presented here that locate *Marronus* within the genus *Oryctes*, we revisited the morphological evidence that placed *M. borbonicus* in its own monotypic genus. *Marronus* has previously been classified in the tribe Pentodontini, separated from the Oryctini by the form of the apex of the metatibiae (truncate vs. toothed, respectively) and by the more pronounced sexual dimorphism of oryctines^27^. However, these characters doubtlessly have evolved numerous times independently and are not taxonomically robust, leaving the monophyly of the tribes in doubt^28^. Truncate metatibiae are a frequent accompaniment to the suite of characters that are related to flightlessness, e.g. dwarfism, atrophied wings, reduced eyes, and thickened legs. Other island flightless rhinoceros beetles purported to be genera of Pentodontini such as *Neoryctes* Arrow (Galapagos Islands) and *Mellissius* Wollaston (St. Helena) display truncate metatibiae, as do numerous other flightless scarabaeoid beetles^29^. Most importantly, *Marronus* displays the same sexually dimorphic characters found in *Oryctes* species, just in a reduced form (Figure 4). These consist of a frontal horn and distinct anterior pronotal fovea in males and a frontal tubercle and smaller pronotal fovea in females. The pronotal fovea of male *Oryctes* is often bordered behind by a bifurcate process and laterally by coarse punctures, and these structures can also be found in males of *Marronus* in reduced form. Thus, the morphology of *Marronus* is consistent with that of *Oryctes* species, agreeing with our phylogenomic analyses, and therefore, *Marronus* should be transferred to *Oryctes*. As two species of the same genus cannot retain identical names, a replacement name will be designated elsewhere.

**Figure 4.**
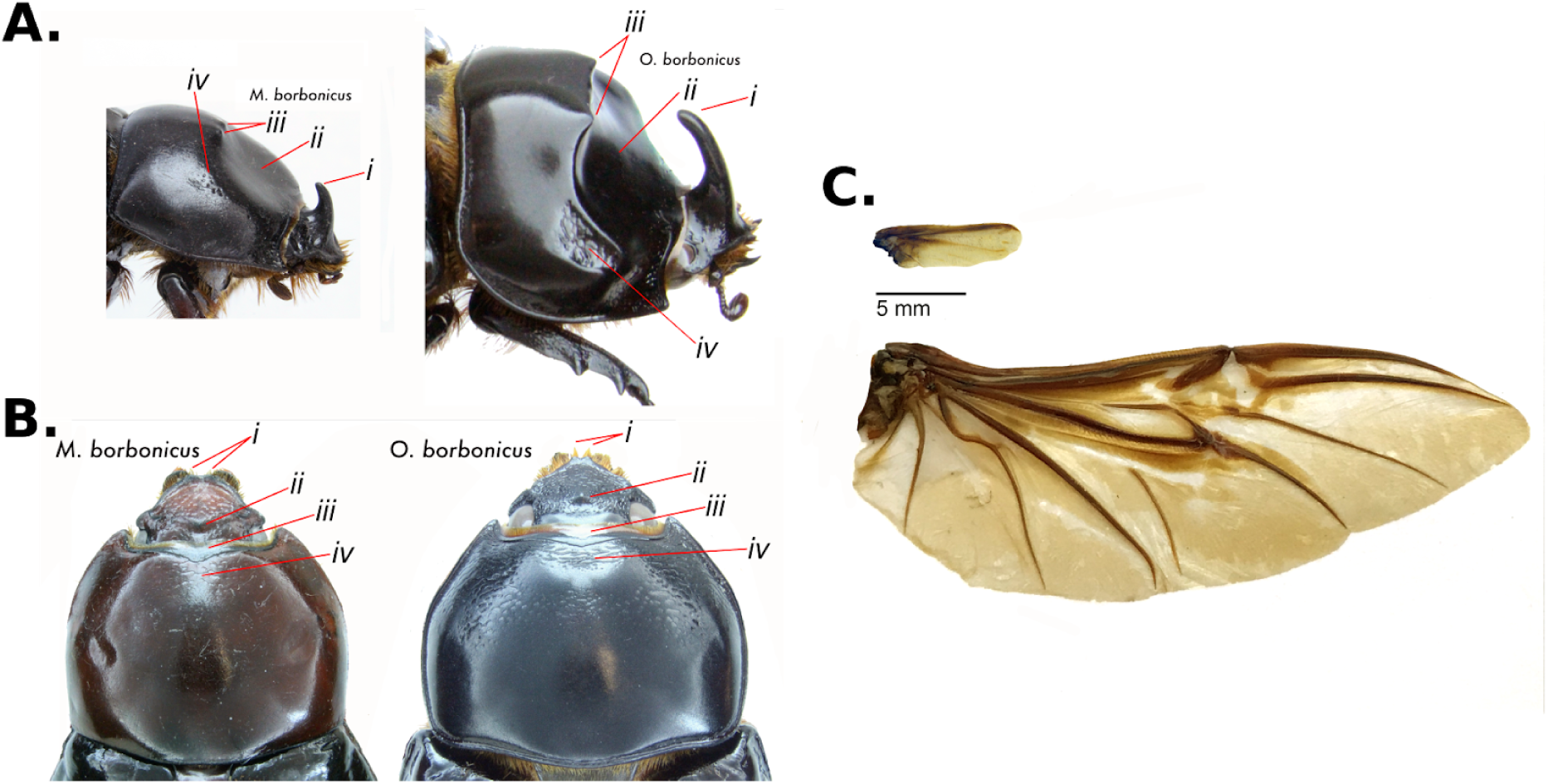
Morphological analysis of *Oryctes borbonicus* and *Marronus borbonicus*. **(A)** Similarities in male specimens of *M. borbonicus* (left) and *O. borbonicus* (right). Roman numbers and red lines point to the presence of analogous morphological characters: *i)* horn, *ii)* pronotal fovea, *iii)* bifurcate process, and *iv)* coarse punctures. **(B)** Similarities in female specimens of *M. borbonicus* (left) and *O. borbonicus* (right). Roman numbers and red lines point the presence of analogous morphological characters: *i*) bidentate clypeus, *ii)* central tubercle, *iii)* anterior margin of pronotum thickened medially, and *iv*) flattened area anteromedially with transverse rugae. **(C)** Wings of *M. borbonicus* (upper) and *O. borbonicus* (lower). While the *O. borbonicus* wing shows venation, size and shape typical of a functional beetle wing, *M. borbonicus* wing size and venation are highly reduced.

## Discussion

Current studies of biodiversity are challenged by rapid environmental change driven by factors such as increased pollution, widespread habitat loss and fragmentation, and unprecedented climate change, which has resulted in increasing extinction rates^30^. Island ecosystems are particularly vulnerable to these threats due to their isolation and ecological uniqueness, which is characterized by high levels of endemism. It is estimated that 80% of species extinctions have taken place in islands and 45% of Red List endangered species also inhabit islands. Thus, while characterizing biodiversity and the evolutionary relationships among different groups of organisms is today more relevant than ever, species are vanishing from Earth at a higher rate than our capacity to study them. Hence, it is becoming necessary to take a retrospective approach to the study of biodiversity that resides in the global archive provided by natural history collections^31^.

Here, we took such a retrospective approach to study the phylogenetic relationships among extant and extinct beetles from the Mascarene Islands. In contrast to most aDNA studies, which usually investigate species that went extinct thousands (e.g. mammoths^32^) or hundreds of years ago (e.g. moas^33^), we studied species that went extinct only a few decades ago. Comparisons between recent extinctions and closely-related extant relatives will shed light on the genetic and ecological factors driving either extinction or persistence of species. To generate genome-wide datasets from historical pinned insect museum specimens, we used state-of-the-art ancient genomics methods including the use of preventive measures to avoid contamination by exogenous DNA, an extraction protocol that enriches for short DNA fragments^18^, and library preparation methods tailored for aDNA^20,34^. The use of these methods permits responsible analysis of precious historical samples with minimal tissue destruction and to present positive evidence of the historical nature of the sequenced DNA^35^.

Our phylogenomic analyses support two independent colonizations of rhinoceros beetles to Réunion island and suggest that *Marronus* became flightless *in situ*. Importantly, estimated sequence divergence times are in agreement with the well accepted geological age of Réunion. Réunion is believed to have emerged about 3 mya and several biological reasons suggest that it has been colonized after Mauritius and Rodrigues, i.e. the absence of flightless birds only on Réunion. In contrast to Réunion, our estimated sequence divergence time for *O. minor* from Rodrigues disagrees with the generally accepted geological age of this island. Specifically, our data suggest that *O. minor* emerged around 5 mya, a finding that is consistent with data on other biota on the island^36^. However, geological studies of Rodrigues are contradictory with more recent work identifying older relicts of lava that supports an age similar to that of Mauritius^36^. These recent findings are more in agreement with our estimated divergence times. Thus, phylogenomics combining extant and extinct species can provide important biological support for the geology and colonization of islands with limited or contrasting geological data.

Our study showcases an integrative taxonomic approach that combines traditional morphological analyses with genome-wide variation from extant and extinct species. In light of current global environmental challenges, and given the vast number of plant and animal collections curated in natural history collections, the widespread use of this approach will be fundamental to catalogue Earth’s biodiversity through space and at different timescales.

## Acknowledgements

We would like to thank R. Gutaker for laboratory assistance, including the implementation of the DNA extraction protocol; E. Reiter for access to clean-room facilities and technical support; Vrinda Venu for technical support during linked-read library preparations; F. Jones for useful input during linked-read the library preparations and genome assembly; K. Pruefer, A. Andres and members of the Burbano and Sommer laboratories for useful discussions and input on data analysis; D.Weigel for supporting S. Latorre’s visit to London during the final stages of the project; and A. Andres, R. Gutaker, T. Karasov and M. Werner for comments on the manuscript. We are also indebted to the Parc national de la Réunion and the Office National des Forets on Réunion for long-term support and permits; the CIRAD at St. Pierre de la Réunion for housing the Max Planck laboratory; J. Rochat and Micropoda for entomological expertise and logistic support; and Max Barclay from the Natural History Museum London and G. Cuccodoro from the Natural History Museum Geneva for providing historical museum samples.

## Author Contributions

S.M.L, R.J.S and H.A.B. conceived and designed the study with input from M.H. and C.R.; S.M.L. carried out DNA extractions and library preparations of historical museum specimens; W.R. and A.D. performed linked-read library preparations; A.D. and C.R. carried out *de novo* assembly of extant genomes; S.M.L. performed phylogenomic and population history analyses with input from H.A.B.; M.H. and M.J.P. performed morphological analyses and contributed entomological expertise; S.M.L., M.H., M.J.P., C.R., R.J.S and H.A.B. contributed to the interpretation of the data; S.M.L., R.J.S and H.A.B. led the writing of the manuscript with input from all authors. All authors read and approved the manuscript.

## Methods

### Fresh samples methods

#### Biological Material

Samples from *Marronus* and *O. borbonicus* were collected in Réunion.

#### DNA Extraction

In order to recover DNA from beetle specimens, two legs plus one thoracic muscle were manually extracted by tweezers from one male specimen of an *O. borbonicus* beetle and three legs plus one thoracic muscle for *Marronus*. The beetle material was ground in liquid nitrogen and processed further according to the QIAGEN Genomic DNA Kit using the 100/G Genomic-tips. DNA was precipitated with 7µl of GlycoBlue (Invitrogen, 15mg/ml) and resuspended in 35µl of EB Buffer (Qiagen). DNA quality was checked by Nanodrop, Qubit and Pulse Field.

#### Library preparation, sequencing and assembly

The preparation of the linked-read sequencing library was done as described previously^37^. Raw sequences were assembled into draft assemblies with the help of *SuperNova*^*38*^ (v. 2.0.1). *SuperNova* was run on the full sequencing data set as well as multiple downsampled read sets. In terms of assembly contiguity, the best results were obtained using around 170 million single reads, which translates into roughly 60X coverage per genome. Assembly quality was assessed using an approach based on benchmarking of universal single copy orthologs (BUSCO^39^). The raw genome assemblies as well as a data set of 1658 orthologous genes from 42 insect species (insecta_odb9) were taken as input of the program *run_BUSCO.py* (v. 3.0.1). To analyse the coverage profile of the previous and current *O. borbonicus* assemblies, we aligned raw reads to both assemblies with the help of the *mem* program of the *BWA* software suite (v. 0.7.17, default options)^40^. Coverage profiles were calculated by the *samtools depth* program (v. 0.1.18, default options)^41^.

#### Gene annotation and comparative genomic analysis

To guide gene annotation, we generated a transcriptome of a male *Marronus* individual by grinding several legs in liquid nitrogen and used the Zymo Direct-zol RNA Miniprep Kit according to the manufacturer’s instructions to extract RNA, which was then eluted in 25µl dH20. We followed previously described methods^42^ to prepare an RNA-seq library and sequenced it on a multiplexed run on a Illumina HiSeq 3000 resulting in 24 million paired end reads (2 × 150bp). Transcriptomic data for *Marronus* and reads from a transcriptomic library of *O. borbonicus*^*12*^ were assembled by the software *Trinity* (v. 2.2.0, default settings)^43^. Full or partial open reading frames were called as described previously^42^. In cases where *Trinity* annotated multiple isoforms, the isoform with the longest ORF was chosen as a representative isoform for subsequent analysis. The resulting ORFs and protein sequences were mapped against their reference assemblies by the *protein2genome* program of the *exonerate* package (version 2.2.0, --bestn 1 --dnawordlen 20 --maxintron 20000)^44^. Among all gene annotations which share an identical exon, the annotation resulting in the longest protein product was taken as representative annotation. Pairwise BLASTP (v. 2.6.0, e-value 0.00001) searches were done between proteins of *Marronus* and *O. borbonicus* and best reciprocal hits were extracted to estimate the median protein sequence identity between orthologous gene pairs. For further comparative genomic analyses, we obtained protein sequences for *Nicrophorus vespilloides* and *Aethina tumida* from the i5k website (https://i5k.nal.usda.gov/, accessed July, 12th 2019)^45^, *Anoplophora glabripennis, Dendroctonus ponderosae, Tribolium castaneum* from Ensembl Metazoa (release 44), *Protaetia brevitarsis, Pyrocoelia pectoralis, Hycleus cichorii* and *Hycleus phaleratus* from GigaDB^46–48^, *Onthophagus taurus* from the U.S. Department of Agriculture website (https://data.nal.usda.gov, accessed July, 12th 2019), *Agrilus planipennis* and *Leptinotarsa decemlineata*^49^ from the ftp server of the Human Genome Sequencing Center of the Baylor College of Medicine (ftp://ftp.hgsc.bcm.tmc.edu/I5K-pilot/, accessed July, 12th 2019), and *Hypothenemus hampei* from the website of the NYU Center for Health Informatics and Bioinformatics (https://genome.med.nyu.edu/) ^50^. In cases of multiple isoforms per gene, the sequence corresponding to the longest protein was taken for further analysis. Assignment of protein sequences into orthologous clusters, concatenation, and phylogenetic reconstruction were performed as described previously^42^.

### Museum samples methods

#### Biological Material

Museum samples provenance is provided in Supplementary Table 2.

#### DNA Extraction

To prevent contamination by exogenous DNA, museum samples were handled using standard ancient DNA precaution measures, i.e. sterilization with UV light of all equipment, surfaces and hoods after each extraction round, and the use of different hoods for handling of samples, reagents and DNA extracts, and of protective gear by researchers. DNA extractions were carried out in the clean-room facility at the Institute of Archeological Sciences at the University of Tübingen. The tissue (one leg per specimen) was ground inside a microtube with a stainless steel pestle until finely powdered and a N-phenacylthiazolium bromide (PTB) and Qiagen Plant DNEasy® Mini Kit (Qiagen)-based protocol was used to isolate the DNA^18^. A microtube without tissue was used as negative DNA extraction control.

#### Library preparation and sequencing

Genomic libraries for all museum specimens were prepared in a DNA clean-room facility taking the same preventive measures described in the DNA extraction section.

#### Non-UDG treated library preparation

We used a protocol that permits the preparation of indexed sequencing ancient DNA libraries^34^. After adapter ligation in the clear-room facility, the indexing and PCR amplification of the libraries libraries were performed in a different laboratory, located in a separate building. Briefly, the libraries were indexed using two barcoded primers^51^ during 10 cycles of PCR amplification using AccuPrime™ Pfx polymerase (Thermo Fisher Scientific). The MinElute PCR Purification Kit (Qiagen) was used to clean PCR residues and samples were pooled in equimolar concentrations. The samples were sequenced at the Genome Center facility located at the Max Planck Institute for Developmental Biology, with the Illumina MiSeq Platform using MiSeq Reagent Kit v2, 300 cycles (Illumina). Together with the prepared libraries, both aDNA extraction and a library preparation negative controls were sequenced. All non-UDG library sequences were used only for authentication purposes and not included in any further analyses.

#### UDG treated library preparation

In spite of the fact that cytosine to thymine (C-to-T) substitutions are useful for the authentication of the samples, they are not desirable in the phylogenomic inferences^19,52^. Thus, in order to reduce the effect of C-to-T substitutions, we prepared new DNAlibraries adding uracil-DNA glycosylase (USER™ enzyme (New England Biolabs)) during the blunting step^20^. The rest of the steps were done as described in the non-UDG treated library section. Finally, after measuring the final DNA concentration per sample and assessing the DNA endogenous concentration (Supplementary Figure 3H), we prepared a pool with a calculated equimolar content of endogenous molecules. Sequencing was done using the Illumina HiSeq 3000 platform (Illumina) located at the Genome Center facility at the Max Planck Institute for Developmental Biology.

#### Pre-processing of reads

Raw reads were trimmed and merged with *AdapterRemoval* (v. 2.2.2)^53^. Merged reads were aligned to the *O. borbonicus* and *M. borbonicus* assemblies presented here using *bwa mem* (v. 0.7.12)^40^. Subsequently, PCR duplicates were removed with *picard tools* (v. 2.8.1).

#### Museum specimens authentication

Ancient DNA (aDNA) has multiple signatures which can be used to its authentication: i) C-to-T substitutions are expected to be enriched at the ends of reads with a decay towards the inner part of the molecule, ii) aDNA is expected to have shorter fragments than fresh DNA molecules and, iii) aDNA is expected to be a mixture between endogenous DNA and non-endogenous DNA^54,55^. We used *MapDamage* (v. 2.0)^56^ to analyse both the C-to-T misincorporation patterns and fragment lengths. We also calculated the endogenous DNA content by dividing the number of mapped reads by the total number of reads. All museum samples displayed the expected signatures of museum / historical samples (Supplementary Figure 3).

#### Identification of segregating positions

To achieve a comparable coverage between historical and fresh samples, four million raw reads from both fresh *O. borbonicus* and *Marronus* were subsampled with the program *samtools view* (v. 1.4)^41^. Together with the museum mapped reads, we created a single bam file with the program *samtools merge* (v. 1.4)^*41*^ and used it as the input for the discovery of variant positions using the program *bsh-denovo*^*57*^. Only positions with full information were considered (flag *-m = 1*) and the alternative allele was randomly sampled (flag *-a = 0.001*). To account for the effect of the Minimum Allele Frequency (MAF) in the evolutionary relationships, we filtered the positions with both 1/7 and 2/7. Since a MAF of 1/7 favours variant sites privative to the more genetically distinct individuals, we used a MAF of 2/7 for all downstream analyses (Supplementary Figures 5 and 6).

#### Phylogenetic analysis

A matrix of pairwise Hamming distances between the individuals was calculated using *Plink* (v. 1.9)^58^ and PCA was computed with the function *prcomp* from the *R* package *stats* V. 3.4.4^59^. Neighbor-net analysis and NJ tree calculations were performed using *SplitsTree* (v. 4.14.6)^*60*^. To test the phylogenetic relations among the species, we used *D*-statistics^21,22^. Based on the results of the previous analysis, we fixed *Oryctes mayottensis* as outgroup species while testing for the relations between *Marronus* and the rest of the beetles denoted as A and B in the following configuration: *D*(((A, B), *Marronus*), *O. mayottensis*). *D*-statistics were calculated using *popstats*^*61*^. Finally, we formally assessed the phylogeny of the scarab beetles by both a Maximum Likelihood and Bayesian-based methods. *RAxML-NG* (v. 0.9.0)^62^ was used for the reconstruction of a ML-based phylogeny. We chose a GTR+G4 substitution model^63^ and performed 200 bootstrap replicates. We also performed a Markov-Chain-Monte-Carlo-based phylogenetic reconstruction using *BEAST* (v. 2.6)^64^. To reduce the complexity of the model we chose a Strict Clock and to reduce the effect of demographic history assumptions we chose a Coalescent Extended Bayesian Skyline approach^65^. Both logs and trees from four independent MCMC chains of 10 million each, with ESS values over 200, were merged using *LogCombiner*. Finally, the Maximum Clade Credibility Tree was computed using *TreeAnnotator.*

In order to discard biases due to the effect of the chosen reference genome, we used the mapped reads to the *Marronus* reference genome (here presented) and repeated all the analysis following the same methodologies we described here (Supplementary Figure 8).

#### Divergence and time estimation

We calculated nucleotide divergence from each individual against the *O. borbonicus* reference genome. The values were calculated within non-overlapping widows of 100kb as Number of variant positions / Total number of positions. Only bases with a sequence support of >= 3X were selected. Nucleotide divergence were converted into time estimates using the reported estimates for arthropods^23^.

## Data availability

Both sequencing raw data and genome assemblies for *O. borbonicus* and *M. borbonicus* have been uploaded to the European Nucleotide Archive (ENA) under the study accession number PRJEB34604. Museum specimens raw data for both UDG and non-UDG treated libraries, have been uploaded under the study accession number PRJEB36751 (Supplementary Table 2). Pipelines and scripts are available at: https://gitlab.com/smlatorreo/museum_phylogenomics_extinct_oryctes_beetles

## Supplementary Information

**Supplementary Figure 1.**
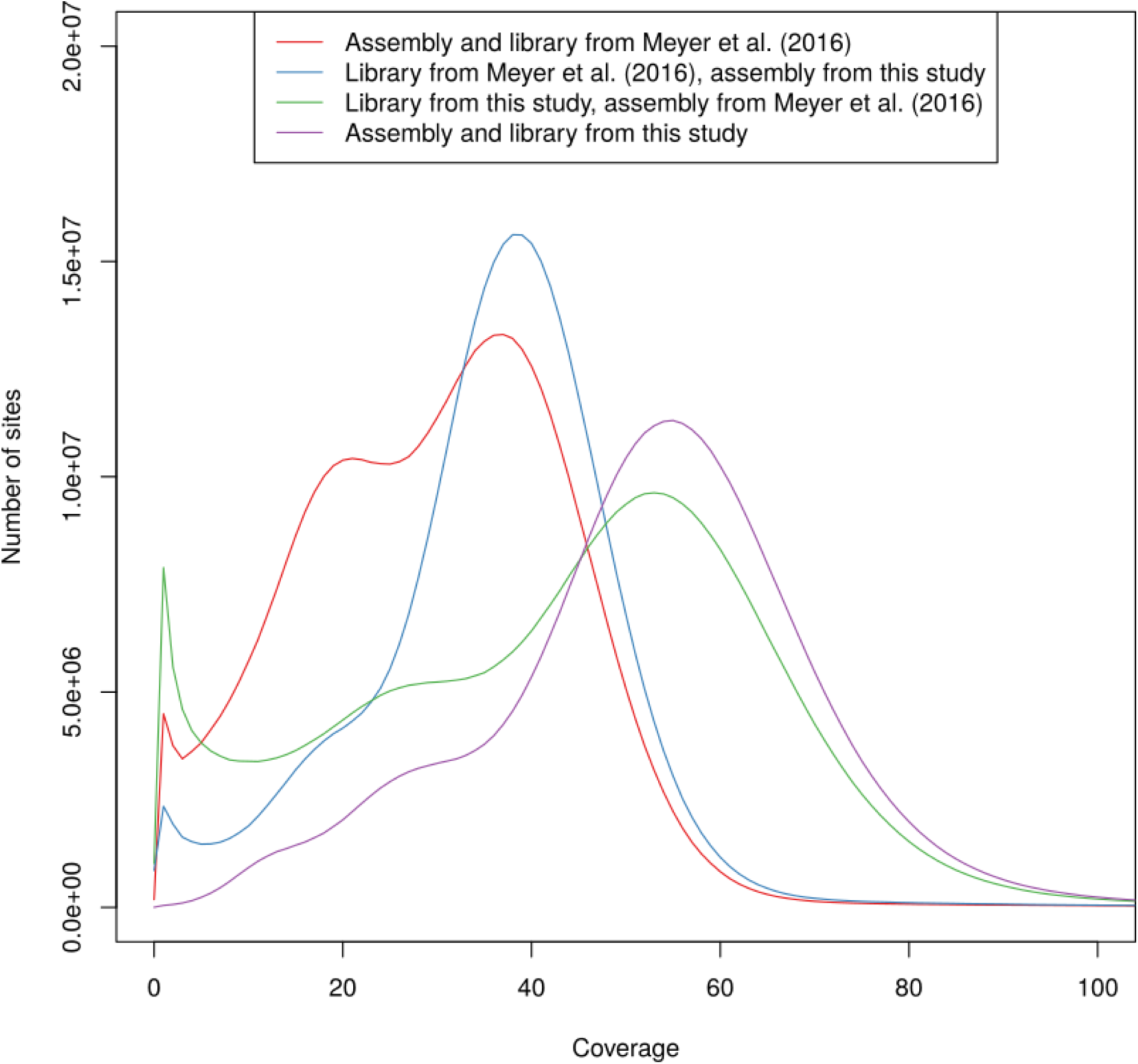
Coverage analysis of different *Oryctes borbonicus* assemblies. Raw reads from two different whole genome sequencing libraries were aligned to the current and the previously published *O. borbonicus* assembly ^12^. For both data sets, the previously published assembly shows a more pronounced peak at half of the expected coverage pointing at the potential problem of allelism in the old assembly.

**Supplementary Figure 2.**
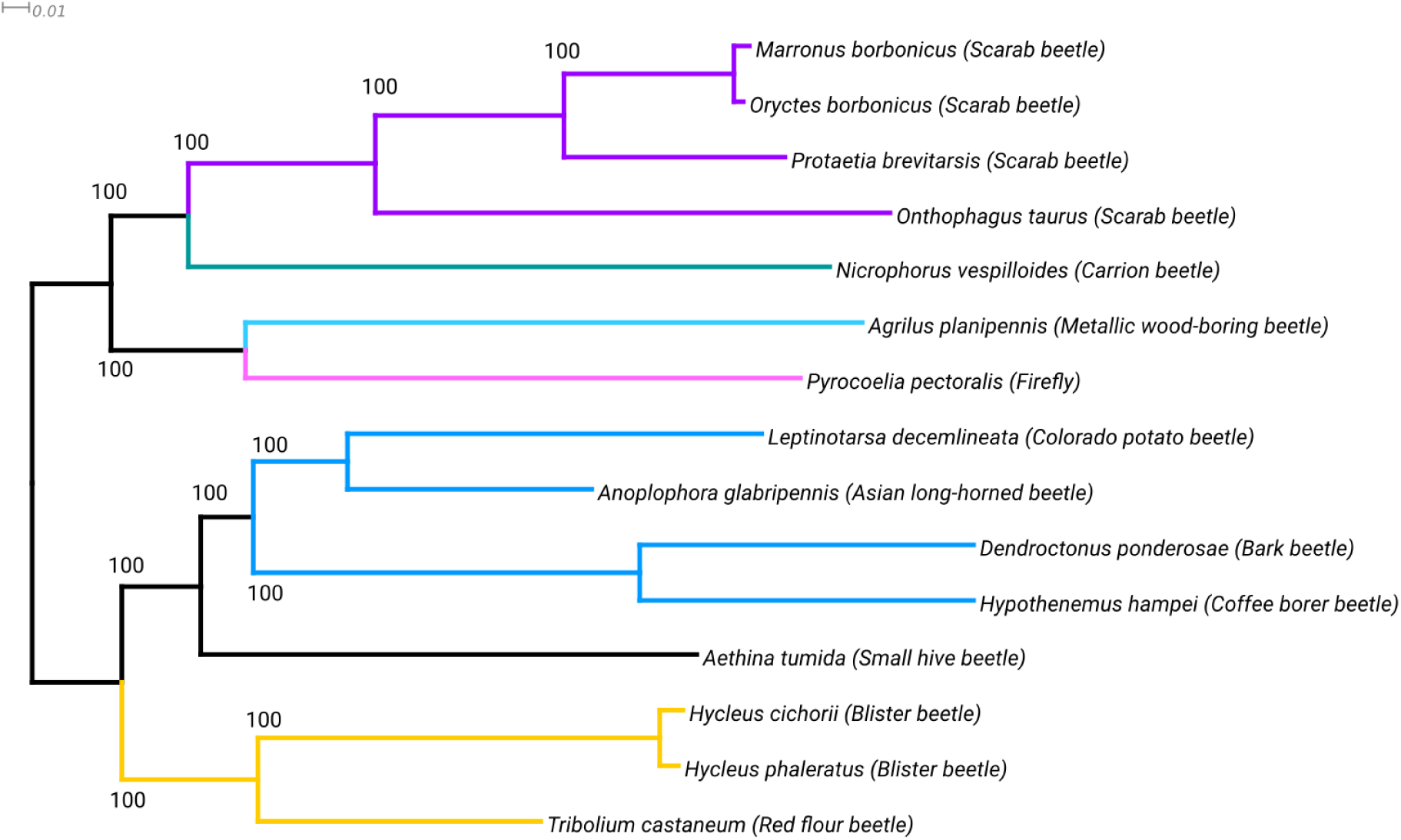
Phylogenetic tree based on genome data from 15 beetle species. Protein sequences from 363 orthologous genes were concatenated. A Maximum likelihood tree was calculated based on the resulting alignment of 107,398 amino acids (100 bootstrap pseudoreplicates). Subtree coloring was chosen for easier comparison with the phylogeny by McKenna et al. ^11^. The two focal species, *Oryctes borbonicus* and *Marronus borbonicus*, display similar levels of divergence as two species of the same genera, *Hycleus cichorii* and *H. phaleratus*.

**Supplementary Figure 3.**
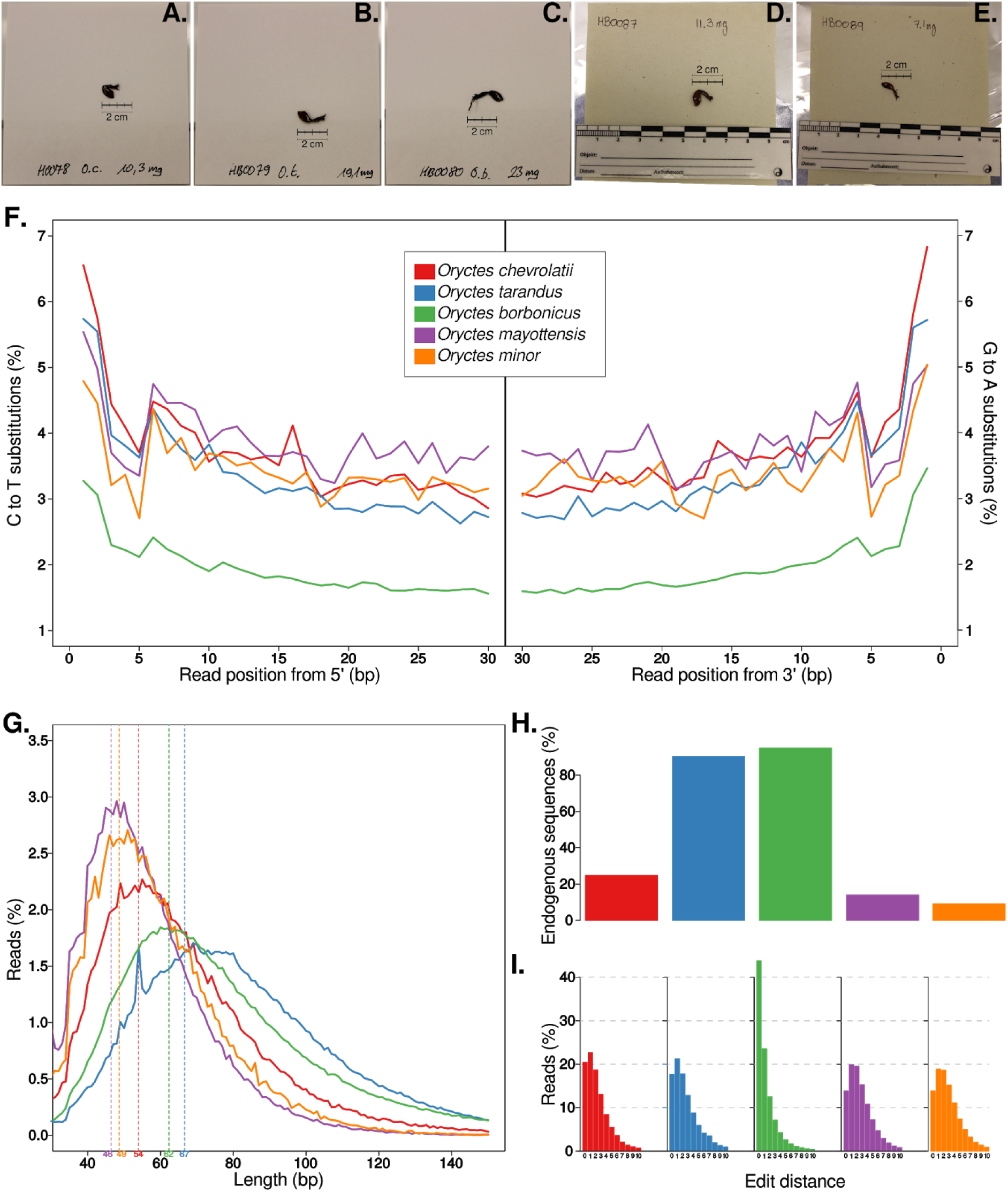
Ancient DNA characteristics and edit distances of museum specimens. **(A-E)** Pictures of samples used for the DNA extraction: **(A)** *Oryctes chevrolatii*, **(B)** *O. tarandus*, **(C)** *O. borbonicus*, **(D)** *O. mayottensis*, **(E)** *O. minor*. **(F)** Cytosine to Thymine and Guanine to Adenine substitutions at the 5’- and 3’-end, respectively. **(G)** Distribution of fragment lengths of merged reads. Dotted lines indicate the mean value of each distribution. **(H)** Percentage of merged reads that mapped to the *O. borbonicus* genome. **(I)** Distribution of edit distances of reads mapped to the *O. borbonicus* genome.

**Supplementary Figure 4.**
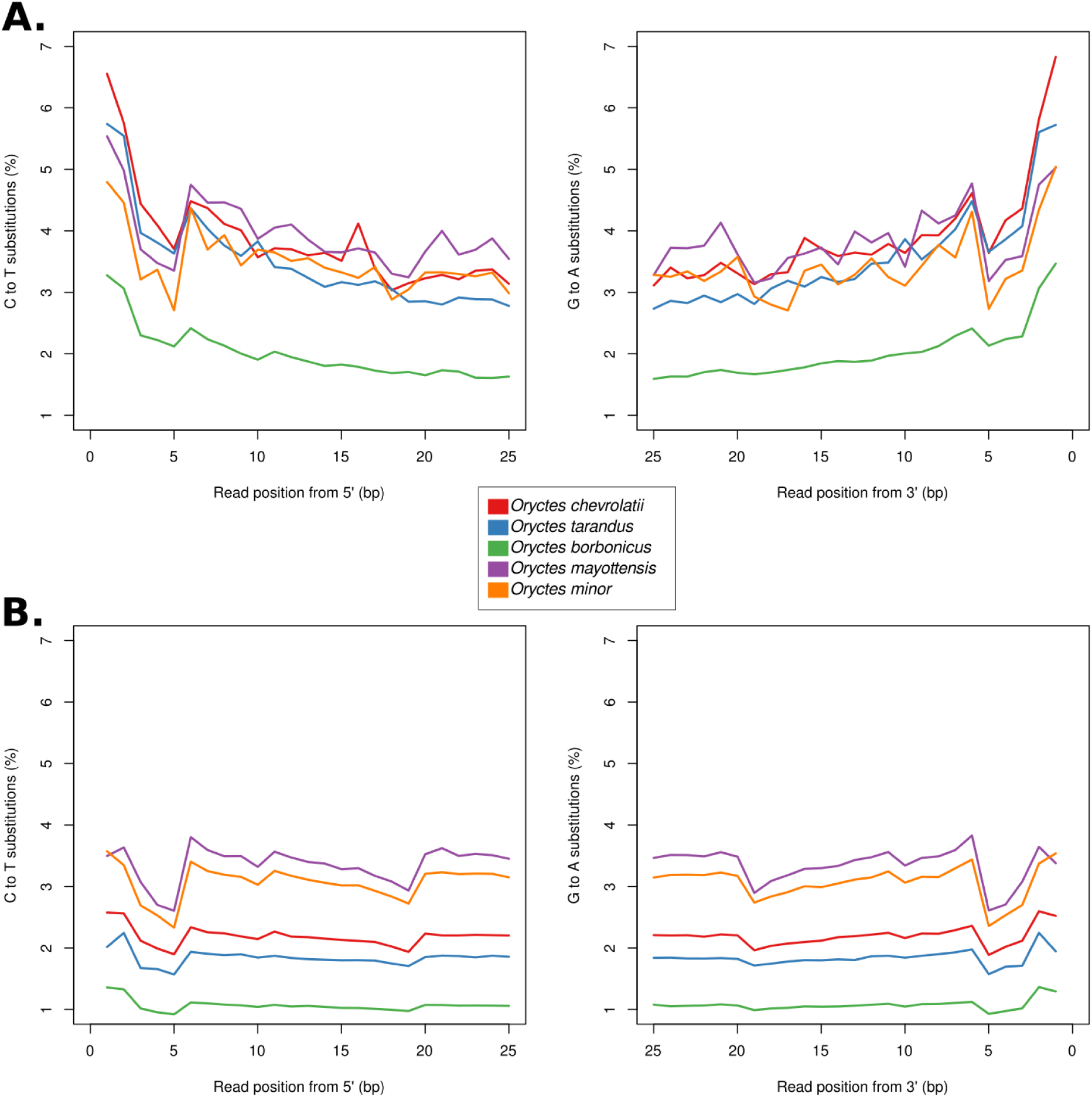
Cytosine to Thymine and Guanine to Adenine substitutions at the 5’- and 3’-end, before and after uracil enzymatic library repair. **(A)** Described substitutions present in museum specimens before enzymatic repair (same as Supp. Fig 3F). **(B)** Described Substitution after enzymatic library reparation.

**Supplementary Figure 5.**
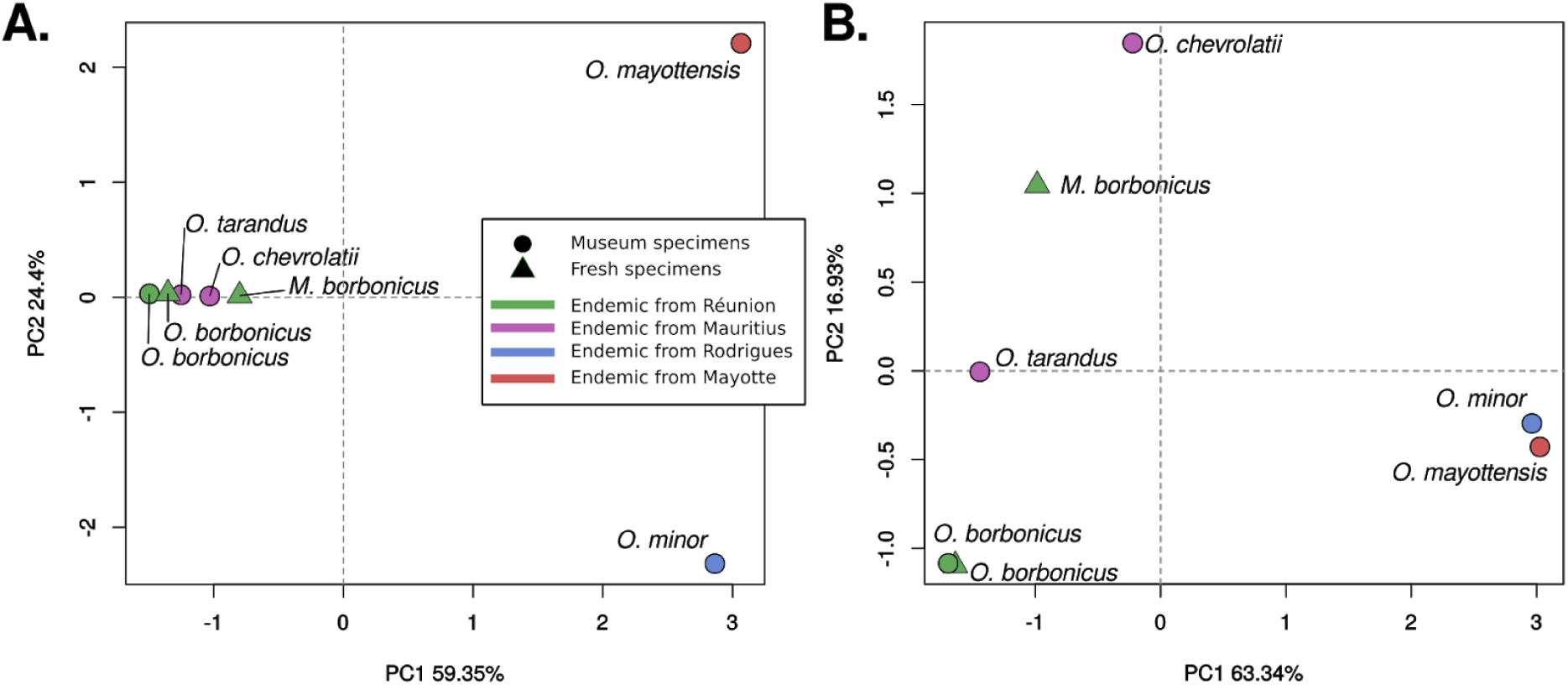
Effect of Minimum Allele Frequency (MAF) filter on the genetic distances. Hamming distances-based PCAs show the effect of the MAF on the separation of the endemic beetles from Réunion and Mauritius. **(A)** MAF of 1/7 with 2,144,289 SNPs and **(B)** MAF of 2/7 with 304,417 SNPs.

**Supplementary Figure 6.**
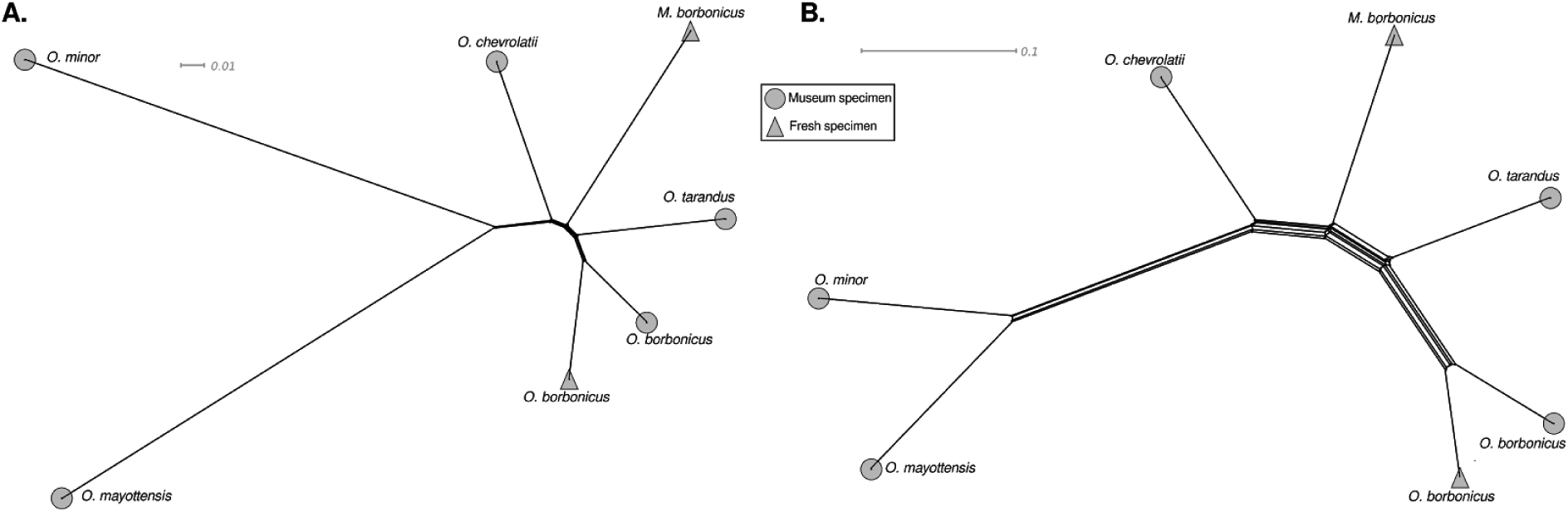
Effect of Minimum Allele Frequency (MAF) filter on the evolutionary relationships and branch lengths. The phylogenetic networks show a similar topology and relations between samples but different branch lengths. **(A)** MAF of 1/7 with 2,144,289 SNPs and **(B)** MAF of 2/7 with 304,417 SNPs.

**Supplementary Figure 7.**
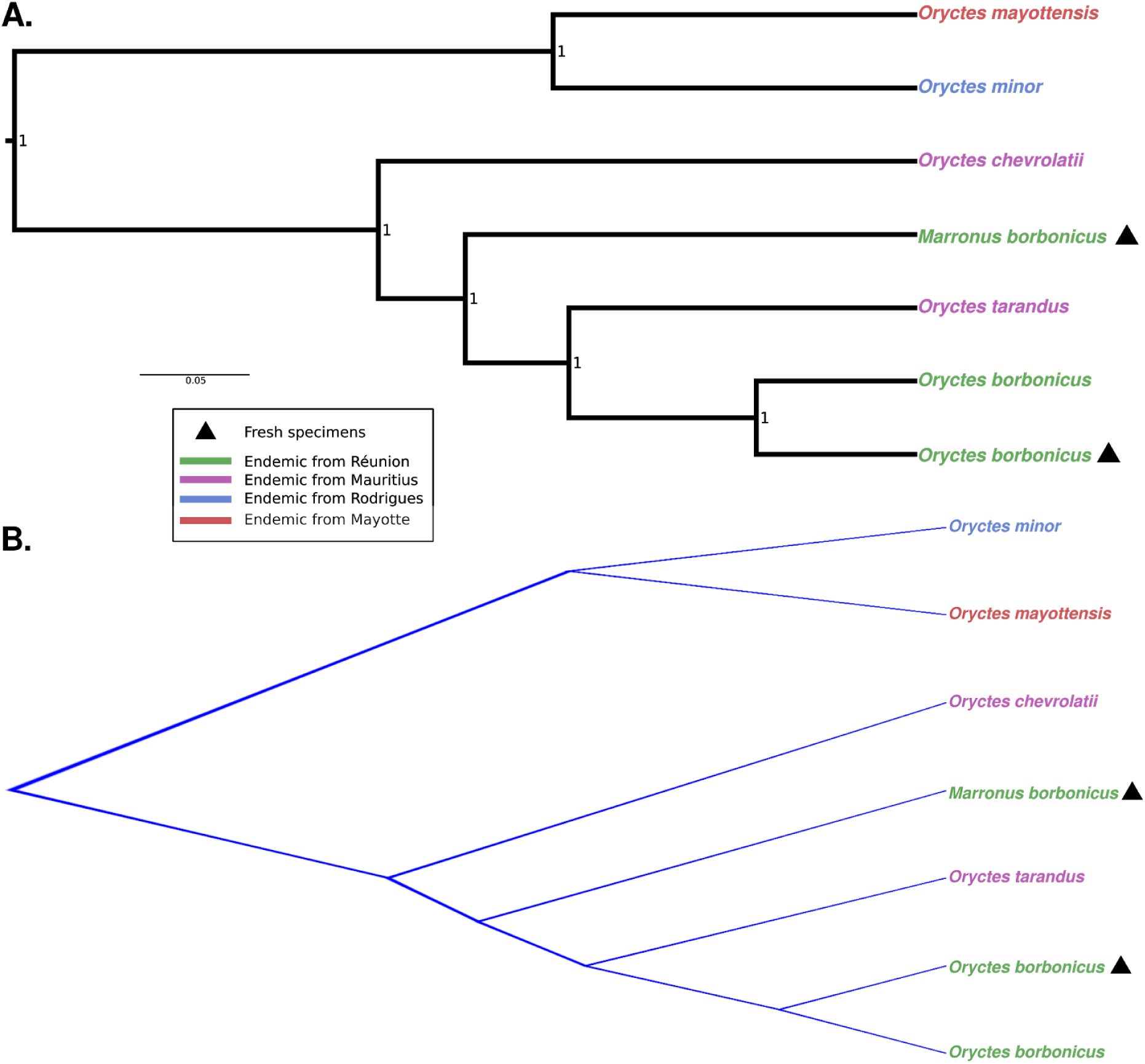
Bayesian phylogenetic tree superposition. **(A)** Bayesian phylogenetic reconstruction. Maximum clade credibility tree. Numbers at the nodes indicate posterior probability support. **(B)** Densitree showing 36,000 trees. In both **A** and **B**, only sites with complete information were included, leaving 304,417 sites in the final dataset.

**Supplementary Figure 8.**
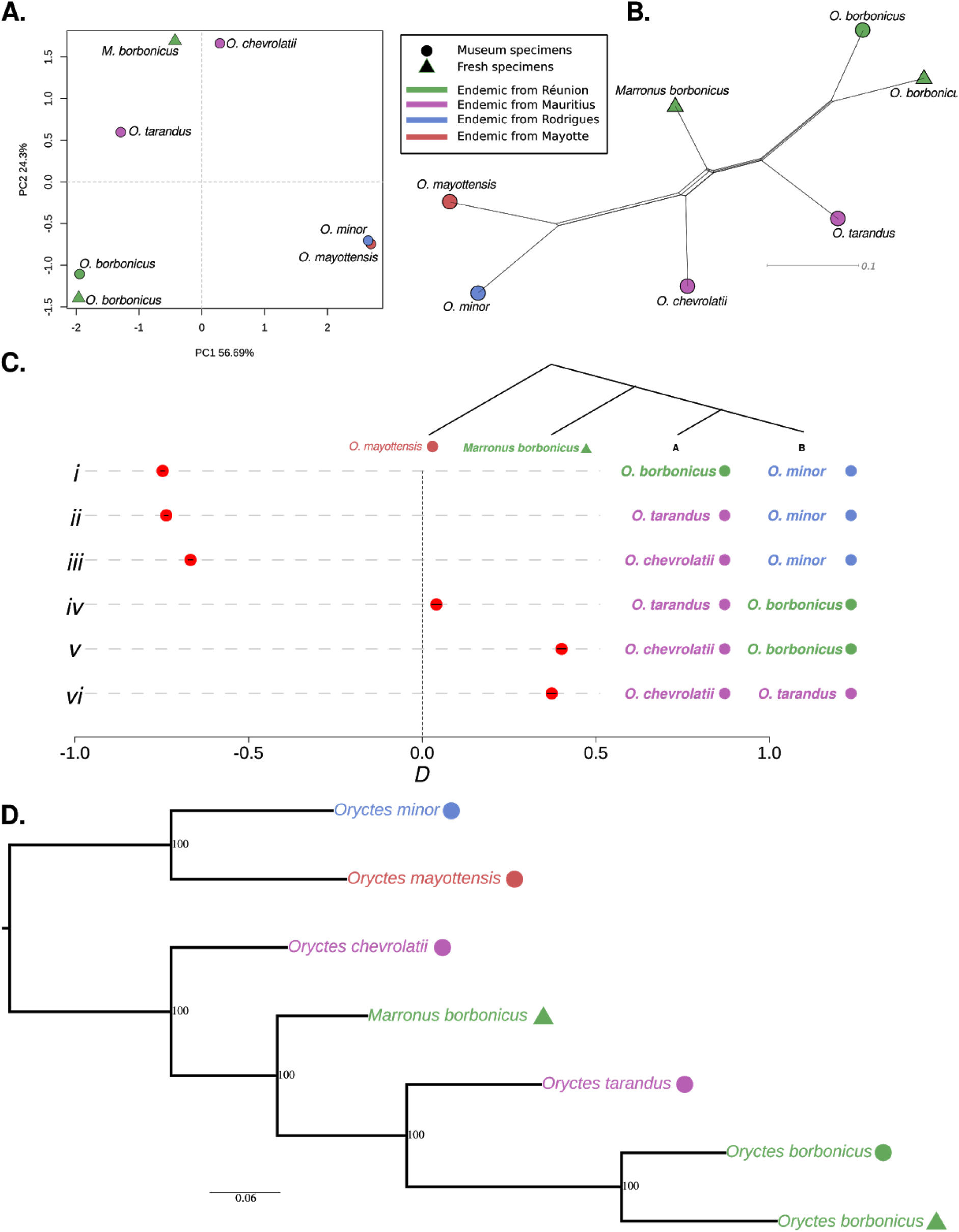
Robustness of evolutionary relations among scarab beetles. The figures represent the same analyses shown in Fig2. In order to discard the possible effect of ascertainment bias due to the selection of *Oryctes borbonicus* as reference genome, we repeated the analysis by mapping the raw reads to a *Marronus borbonicus* assembly. **(A)** Principal component analysis plot based on 330,529 SNPs. Genetic distances between beetle samples are projected onto the first two PCs. Axis labels indicate the fraction of total variation explained by each PC. **(B)** Phylogenetic network based on 330,529 SNPs using the neighbor-net method. **(C)** Testing the robustness of phylogenetic relations among scarab beetle species using *D*-statistics of the type *D*(B,A; *Marronus borbonicus*, outgroup), as depicted in the phylogenetic tree. *O. mayottensis* was used as an outgroup. Each row (*i-vi*) shows a different *D*-statistic configuration. A negative *D*-statistic indicates that *M. borbonicus* is closer to species B, whereas a positive *D*-statistic indicates that *M. borbonicus* is closer to species A. The points depict the result of each *D*-statistic test and the lines their respective 95% confidence intervals. Rows *i-iii* show that *M. borbonicus* is closer to the *Oryctes* spp. from Réunion and Mauritius. Rows *v-vi* show that *M. borbonicus* is closer to both *O. borbonicus* and *O. tarandus* than to *O. chevrolatii*. Finally, row *iv* shows the closest *D*-statistic to zero, which indicates that *M. borbonicus* is slightly closer to *O. borbonicus* than to *O. tarandus*. **(D)** Maximum-Likelihood phylogenetic tree. Numbers at nodes indicate bootstrap support (200 replicates).

**Supplementary Table 1.** Overview of genome sequencing and assembly of extant beetle specimens.

| **Species** | **Ref** | **Number of Scaffolds** | **Genome size (assembled) [Mb]** | **N50 [Mb]** | **BUSCO Complete single copy [%]** |
| --- | --- | --- | --- | --- | --- |
| *O. borbonicus* | (Meyer et al. 2016) | 150,243 | 494.4 | 0.1 | 95.3 |
| *O. borbonicus* | This study | 9,526 | 411.2 | 8.4 | 95.9 |
| *M. borbonicus* | This study | 9,046 | 412.6 | 12.4 | 95.8 |

**Supplementary Table 2.** Samples information.

| ID | (M)useum<br>/ (F)resh | Species | Collection<br>Year | Collection<sup>1</sup> | Collector | Non-UDG<br>treated<br>libraries<sup>2</sup> | UDG treated<br>libraries<sup>2</sup> |
| --- | --- | --- | --- | --- | --- | --- | --- |
| HB0078 | M | *Oryctes chevrolatii* | 1963 | NHMG | Rouillard | ERR3932721 | ERR3932716 |
| HB0079 | M | *Oryctes tarandus* | 1966 | NHMG | Yves Gomy | ERR3932722 | ERR3932717 |
| HB0080 | M | *Oryctes borbonicus* | 1966 | NHMG | Yves Gomy | ERR3932723 | ERR3932718 |
| HB0087 | M | *Oryctes mayottensis* | 2010 | MPI | NA | ERR3932724 | ERR3932719 |
| HB0089 | M | *Oryctes minor* | 1918 | NHML | Snell & Thomasset | ERR3932725 | ERR3932720 |
| - | F | *Oryctes borbonicus* | 2017 | MPI | Matthias Herrmann | ERR3685131<br>-<br>ERR3685134 | NA |
| - | F | *Marronus borbonicus* | 2017 | MPI | Matthias Herrmann | ERR3685135<br>-<br>ERR3685138 | NA |

<sup>1</sup>NHMG (Natural History Museum Geneva); NHML (Natural History Museum London); MPI (Max Planck Institute for Developmental Biology, Tuebingen).
<sup>2</sup> ENA accession numbers.

**Supplementary Table 3.** Average depth in museum specimens mapped to the *Oryctes borbonicus* draft genome.

| Sample | Average depth (X) |
| --- | --- |
| *Oryctes borbonicus* | 1.16 |
| *Oryctes tarandus* | 1 |
| *Oryctes chevrolatii* | 0.79 |
| *Oryctes mayottensis* | 0.8 |
| *Oryctes minor* | 1.38 |

